# PLK1 controls centriole distal appendage formation and centrobin removal via independent pathways

**DOI:** 10.1101/2021.07.06.451279

**Authors:** Morgan Le Roux-Bourdieu, Daniela Harry, Patrick Meraldi

## Abstract

Centrioles are central structural elements of centrosomes and cilia. They originate as daughter centrioles from existing centrioles in S-phase and reach their full functionality with the formation of distal and subdistal appendages two mitoses later. Current models postulate that the centriolar protein centrobin acts as placeholder for distal appendage proteins that must be removed to complete distal appendage formation. Here, we investigated in non-transformed human epithelial cells the mechanisms controlling centrobin removal and its effect on distal appendage formation. We demonstrate that centrobin is removed from older centrioles due to a higher affinity for the newly born daughter centrioles, under the control of the centrosomal kinase Plk1. Centrobin removal also depends on the presence of subdistal appendage proteins on the oldest centriole. It is, however, not required for distal appendage formation even though this process is equally dependent on Plk1. We conclude that during centriole maturation, Plk1 kinase regulates centrobin removal and distal appendage formation via separate pathways.

## Introduction

Centrosomes are the main microtubule nucleating and organizing centres of metazoan cells. Each centrosome is composed of two barrel-shaped centrioles, composed of nine microtubule triplets surrounded by pericentriolar material (PCM) (Gönczy and Hatzopoulos, 2019; Vásquez-Limeta and Loncarek, 2021). Like DNA, centrosomes replicate in a semi-conservative manner once per cell cycle. In G1, cells contain two centrioles: one older centriole called here “grandmother” centriole and one younger daughter centriole. As both give rise orthogonally to one “daughter” centriole during S-phase, the G1 daughter centriole becomes the “mother” centriole. The two new-born daughter centrioles elongate in G2, followed by centrosome separation and maturation at mitotic onset. Therefore, mitotic cells always contain three different centriole generations: one grandmother centriole, one mother centriole and two daughter centrioles.

One key difference between the grandmother centriole and the mother centriole is the presence of distal and sub-distal appendages on the grandmother centriole that emerge as spikes from the centriolar barrel (Tischer et al., 2021). The transition from nascent daughter centriole to grandmother centriole via mother centriole encompasses two mitoses, and one and a half cell cycles (Kong et al., 2014). At the end of the second mitosis, the former mother centriole has recruited the full set of distal and sub-distal appendage proteins, driving the formation of both appendages, and completing to transformation into a grandmother centriole. Each type of appendage ensures key centrosomal or ciliary functions (Loncarek and Bettencourt-Dias, 2018). Sub-distal appendages enable centrioles to anchor and focus microtubules directing intracellular trafficking, cell motility, cell adhesion or cell polarity during interphase (Bornens, 2002; Delgehyr et al., 2005). At its core, sitting on the centriole wall, resides cenexin, which is required for the recruitment of all the proteins that are located in the outer parts of sub-distal appendages, such as CEP128, centriolin or ninein (Chong et al., 2020; Mazo et al., 2016). Distal appendages are required for the hook-like insertion of the older centriole into the plasma membrane during ciliogenesis (Tanos et al., 2013). The binding of OFD1 is the first building step for the formation of distal appendages (Bowler et al., 2019; Wang et al., 2018). This proteins plays a pleiotropic role regulating distal appendage formation, centriole length, and ciliogenesis (Singla et al., 2010). In terms of distal appendage formation, OFD1 binding enables the hierarchical recruitment of all subsequent proteins at mother centrioles, starting with CEP83 and ending with FBF1 and CEP164 (Bowler et al., 2019; Wang et al., 2018). Super-resolution microscopy further revealed that distal and subdistal appendages proteins mutually influence each other’s position relative to the centriole, indicating that they are structurally partially inter-dependent (Chong et al., 2020).

A central player in the current regulatory model of distal appendage formation is centrobin. This protein accumulates in mammalian cells on daughter-centrioles (Zou et al., 2005); it is required for efficient centrosome duplication and essential for cilia formation (Gudi et al., 2014; Ogungbenro et al., 2018). In the context of distal appendage formation, centrobin has been proposed to act as a placeholder for the recruitment of future distal appendage proteins, and its removal under the control of the centriole protein Talpid3 is thought to allow the recruitment of OFD1 (Wang et al., 2018). This raises the question as to the exact timing of centrobin removal during the centrosome cycle, and the nature of the regulatory pathways controlling this process. One likely regulator is the polo-like kinase 1 (PLK1). In Drosophila neuroblasts, which contain appendage-free centrioles, centrobin is exclusively present on the younger centrosome containing the mother centriole, whose identity it controls (Januschke et al., 2013); its removal from the mother centriole in mitosis depends on the PLK1 ortholog Polo (Gallaud et al., 2020). In mammalian cells, PLK1 can phosphorylate centrobin in mitosis (Lee et al., 2010); moreover, its kinase activity is required for the loading of several distal appendage components such as CEP164 (Kong et al., 2014). Therefore, it is thought that PLK1 promotes the formation of distal appendages including the recruitment of CEP164 via the removal of centrobin from the mother centriole. This hypothesis has, however, never been tested in one coherent model system or cell line. Here, we tested in non-malignant, diploid human retina pigment epithelial RPE1 cells the localization pattern of centrobin over the cell cycle, its potential regulation by PLK1, and its role in the formation of distal appendages. Furthermore, we tested how sub-distal appendage proteins contribute to centrobin dynamics, as the sub-distal appendage protein cenexin is a key regulator of PLK1 activity (Colicino et al., 2019). We show that centrobin is removed from the mother centriole as cells reach metaphase, and that this removal depends on the newly emerging daughter centrioles, possibly because of higher affinity binding sites. Centrobin removal and formation of distal appendages depend both on PLK1 activity and the presence of cenexin. Surprisingly, these two centrosome cycle steps occur independently of each other. Indeed, in multiple siRNA or CRISPR/Cas9 knock-out backgrounds, distal appendage formation occurs in RPE1 cells without centrobin removal. We therefore propose that PLK1 and sub-distal appendage proteins regulate the removal of centrobin and the build-up of distal appendages via separate pathways, pointing for a need to revise the regulatory model for centriole maturation.

## RESULTS

### Centrobin is removed from the mother centriole at the prometaphase-metaphase transition

Centrobin was first described as a daughter centriole-specific protein that localizes to the daughter centriole in G1 before appearing on the two new daughter “pro”-centrioles that arise from the grandmother and the new “mother” centriole in S-phase (Zou et al., 2005). How and when centrobin is removed from the mother centriole in human cells is, however, unknown. To address the when, we studied the cell cycle-dependent localization of centrobin in human retina-pigment epithelial cells immortalized with telomerase and expressing the centriole marker eGFP-centrin1 (hTertRPE1-eGFP-centrin1). Cells were fixed for immunofluorescence, stained for centrobin, the S-phase marker PCNA, and the DNA marker DAPI, and assigned to: 1) G1, if the cell was PCNA-negative and contained two centrioles; 2) S-phase, if the cell was PCNA-positive - note that only cells with 4 distinct centrioles were considered; 3) G2 if the cell was PCNA-negative and contained four centrioles. In addition, an increasing abundance of eGFP-centrin1 on daughter, mother and grandmother centrioles allowed to classify each centriole in one of those age category (Tan et al., 2015). In most G1 cells, centrobin was only present on the daughter centriole (91±4%; Fig. 1A and B). In S-phase cells that had initiated centriole duplication and hence contained four eGFP-centrin1 dots, most cells displayed centrobin on both daughter centrioles and the mother centriole (64,5±7%; Fig. 1A and B). Finally, in G2 we observed a mix of three populations: 27,4±5.6% cells with centrobin on all four centrioles, 34.3±4.7% of cells with centrobin on the mother and the two daughter centrioles, and 38.2±3.7% of cells with centrobin only on the two daughter centrioles (Fig 1. A and B). We conclude that centrobin localization is more dynamic than previously estimated, and that centrobin can simultaneously be present on daughter and mother centrioles or even the grand-mother centriole up to G2 phase.

**Figure 1:**
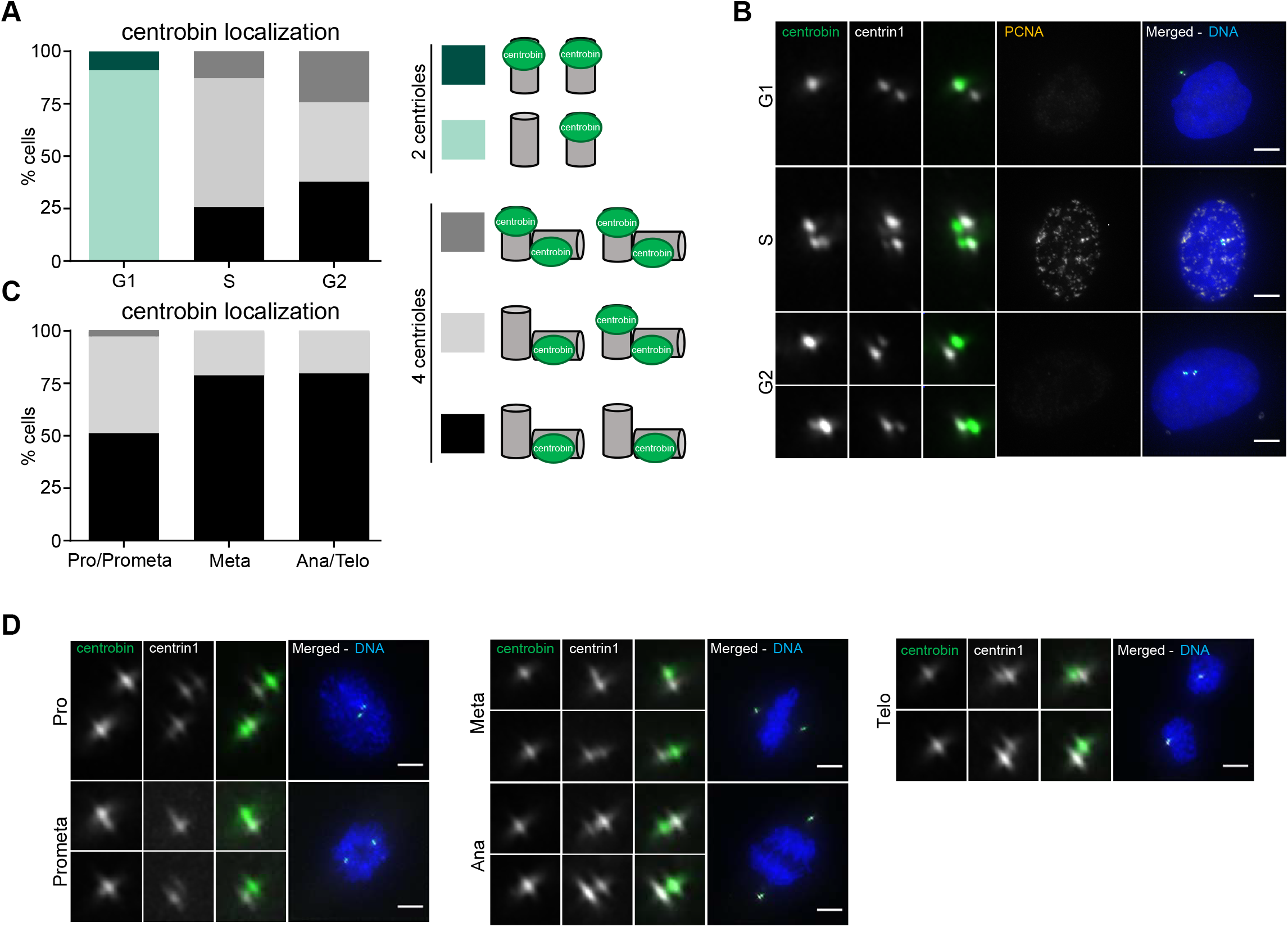
Centrobin is removed from mother centrioles at the prometaphase/metaphase transition. **(A**) Quantification of the centrobin localization pattern in hTert-RPE1-eGFP-centrin1 cells in G1, S or G2. N ≥ 3 independent experiments, n = 31-133 cells. (**B**) Immunofluorescence images of hTert-RPE1-eGFP-centrin1 stained with DAPI and antibodies against centrobin and PCNA as quantified in A. (**C**) Quantification of the centrobin localization pattern in hTert-RPE1-eGFP-centrin1 cells in in the different mitotic phases. N= 2, n = 63 - 74 cells. (**D**) Immunofluorescence images of hTert-RPE1-eGFP-centrin1 stained with antibodies against centrobin and DAPI as quantified in C. All scale bars = 5 μm.

To determine the exact time-point at which centrobin is removed from mother centrioles, we analysed its localization pattern in the different mitotic phases, using DAPI to determine the mitotic stage. In prophase and prometaphase roughly half the cells still displayed centrobin on the daughter centrioles and the mother centriole, while the other half contained only two centrobin dots, on the daughter centrioles (Fig. 1C and D). From metaphase on, 79% of the cells only displayed two centrobin dots, exclusively on the daughter centrioles (Fig. 1C and D). As this percentage did not significantly increase in anaphase and telophase (Fig. 1C and D), we conclude that the majority of centrobin is removed from the mother centriole at the prometaphase/metaphase transition.

### Centrobin is removed from mother centriole due to a higher affinity for daughter centrioles

To better understand the pathways controlling centrobin localization, we next tested whether centrobin removal was linked to the formation of fully elongated daughter centrioles in mitosis. To eliminate daughter centrioles, we inhibited for 18 hours PLK4, the master regulator of centriole duplication, with the chemical inhibitor centrinone (Wong et al., 2015). This resulted in the formation of cells that entered mitosis with only one centriole at each centrosome (called “1:1” cells). When compared to DMSO-treated “2:2” cells or to 2:2 cells that were only treated for 2 hours with centrinone, 1:1 cells formed structurally normal spindle poles in mitosis as measured by the recruitment of γ-tubulin and pericentrin, two of the main components of the pericentriolar matrix (Supplementary Fig. 1A-D). Moreover, these centrosomes were as functional as normal centrosomes with 2 centrioles: 1:1 cells had the same mitotic timing and the same chromosome segregation error rate when released from a prometaphase arrest induced with the centrosome separation/Eg5-inhibitor STLC (DeBonis et al., 2004), as DMSO-treated or 2:2 cells treated with centrinone for only 2 hours (Supplementary Fig. 1E-H). This contrasted with RPE1 cells lacking centrioles on one (1:0 cells) or both poles (0:0 cells), which we and others have found to spend more time in mitosis and have an elevated chromosome segregation error rate (Dudka et al., 2019; Wong et al., 2015).

When we analysed the localization of centrobin in metaphase 1:1 cells, we found that 79.9±6.3% contained a single centrobin-positive centriole (Fig. 2A and B). Using antibodies against the subdistal appendage protein cenexin (grandmother centriole marker; Lange and Gull, 1995), we established that this centriole was as a rule the mother centriole (Fig. 2A and C). The persistence of centrobin on the mother centriole in metaphase was not due to centrinone *per se*, since a short 2-hour centrinone treatment in control 2:2 cells did not maintain centrobin at the mother centriole (Supplementary Fig. 1I). Moreover, inhibition of centrosome duplication by depletion of the centriole duplication seed SAS-6 for 24 hours (Leidel et al., 2005), gave identical outcome to an 18-hour inhibition of PLK4, confirming the specificity of our result (Fig. 2E).

**Figure 2:**
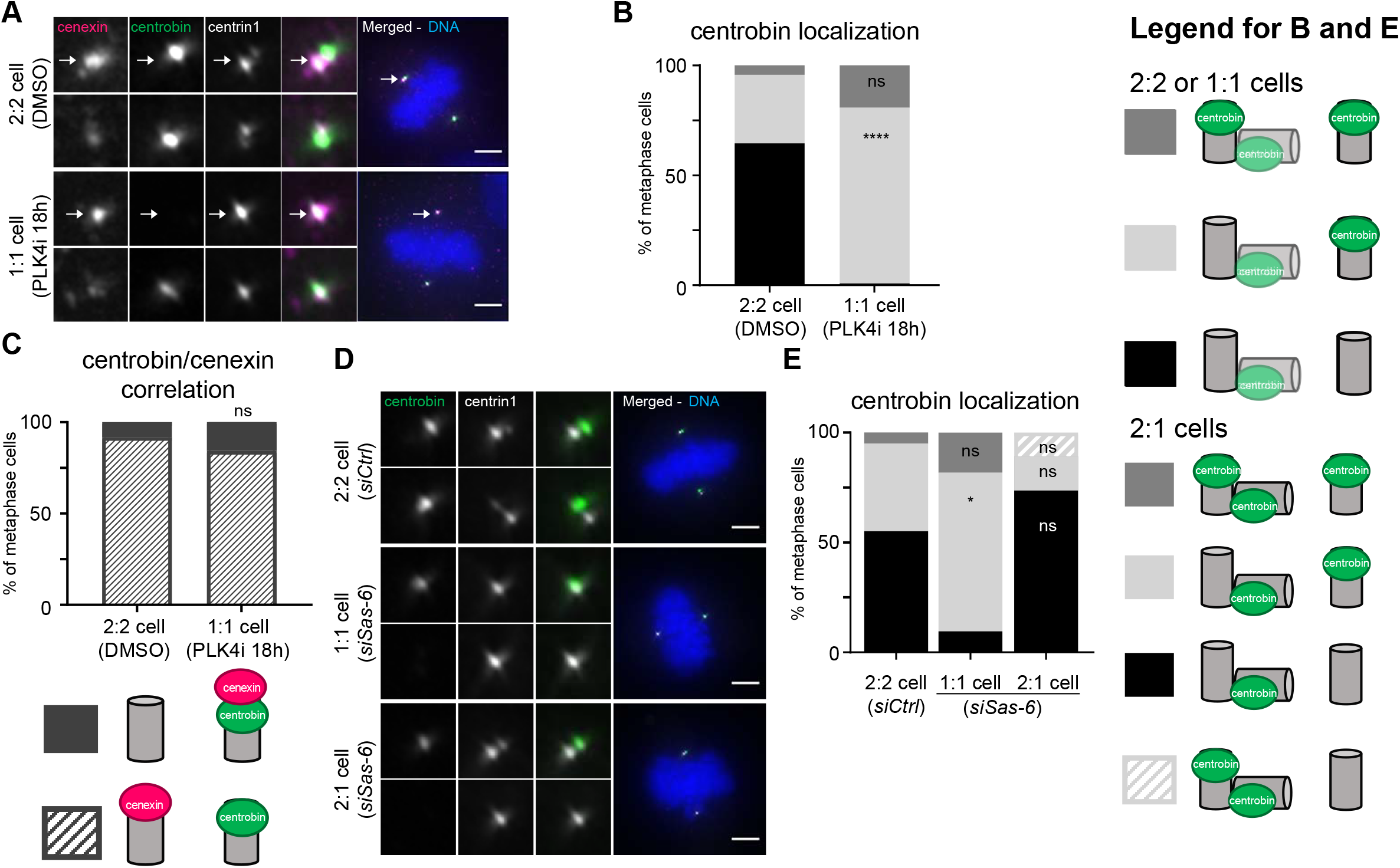
Daughter centrioles compete for centrobin binding with mother centrioles. (**A**) Immunofluorescence images of metaphase hTert-RPE-eGFP-centrin1 cells stained with DAPI and antibodies against cenexin and centrobin after DMSO-treatment or PLK4 inhibition for 18 hours. White arrows identify the older centrosome. (**B**) Quantification of the centrobin localization pattern in hTert-RPE1-eGFP-centrin1 cells treated with DMSO or centrinone for 18h. N = 5, n = 151 - 182 cells, ****p < 0.0001 in two-way Anova (**C**) Quantification of the centrobin/cenexin co-localization in hTertRPE1-1 eGFP-CENP-A/centrin1 after DMSO-treatment or PLK4 inhibition for 18 hours. N = 4, n = 26– 75 cells. (**D**) Immunofluorescence images of metaphase hTertRPE1 eGFP-centrin1 cells treated with *siCtrl* or *siSas-6* for 24h and stained with DAPI and centrobin antibodies. (**E**) Quantification of the centrobin localization pattern in hTert-RPE1-eGFP-centrin1 cells after control or Sas-6 depletion. N = 3 experiments, n = 78 - 121 cells. All scale bars = 5 μm.

The presence of centrobin on the mother centriole in metaphase 1:1 cells had two potential causes: either centrobin removal requires the younger centriole to become a “mother” that gave birth to a daughter centriole, or it is removed due to the presence of a higher affinity site on the fully elongated daughter centrioles. To differentiate between the two possibilities, we looked at the cells depleted for SAS-6 for 24 hours, which in our experience can lead to the formation of 2:1 cells, in which only the grand-mother centriole gives rise to a daughter procentriole (Tan et al., 2015). The vast majority of 2:1 cells contained one centrobin-positive daughter centriole associated to the grandmother centriole, and no centrobin on the mother centriole (Fig 2E). We conclude that centrobin is removed from the mother centriole because daughter centrioles have a higher affinity for this protein.

### Distal appendage proteins are recruited to the mother centriole independently of centrobin removal

Recent studies postulated that distal appendage formation at mother centrioles requires the step-wise removal of the daughter-specific CEP120, centrobin and neurl4 in early S-phase to enable the recruitment of OFD1 (Wang et al., 2018). This first step is followed later at mitotic onset with the recruitment of other distal appendage proteins, including the recruitment of CEP164 (Fig. 3A). Thus, in a normal metaphase cell, OFD1 and CEP164 are present on both the grandmother and mother centriole. To control whether centrobin presence at the mother centriole in 1:1 cells prevented this recruitment, we stained for OFD1: a large majority of 1:1 cells (75±2.5%) still displayed OFD1 at both grandmother and mother centrioles, even though the percentage was slightly lower than in 2:2 cells (91±1.3%; Fig. 3B and C). CEP164 localization was even less affected, as it was present at grandmother and mother centrioles to same extent in 2:2 and 1:1 cells (Fig. 3D and E). We conclude that the presence of centrobin at mother centrioles did not prevent the recruitment of distal appendage proteins implying that centrobin removal and distal appendage recruitment are independent steps of the centrosome cycle. Consistent with this hypothesis, we also found examples of late G2 wild-type 2:2 cells, in which CEP164 and centrobin (albeit weakly) could be found on the same mother centriole (Fig. 3F).

**Figure 3:**
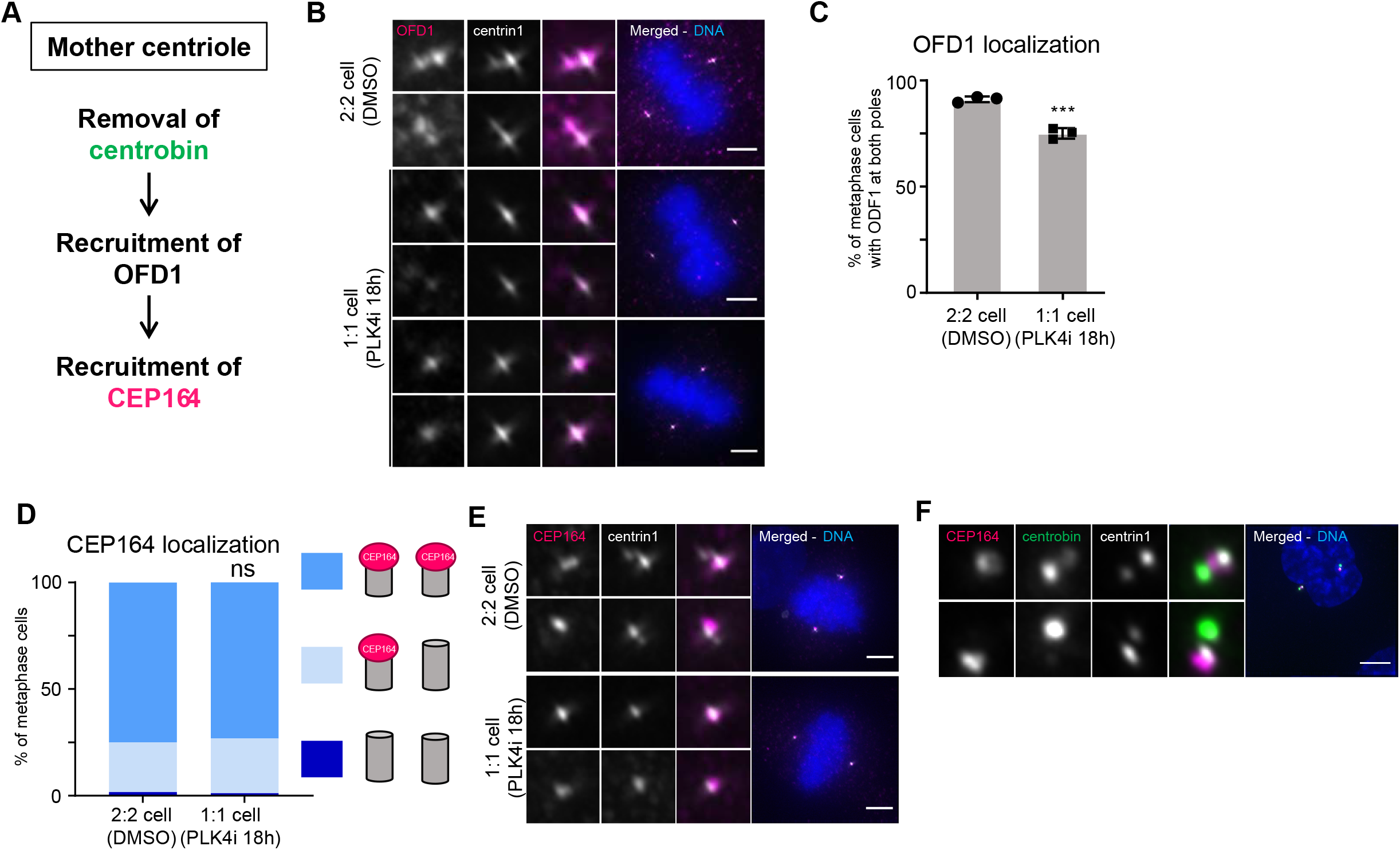
Centrobin removal is not required for the recruitment of distal appendage proteins. (**A**) Model for the pathway controlling CEP164 recruitment on the mother centriole, as proposed by Wang et al., 2018. (**B**) Immunofluorescence images of metaphase hTertRPE1 eGFP-centrin1 stained with DAPI and OFD1 antibodies and treated either with DMSO or centrinone for 18h. (**C**) Quantification of OFD1 localization pattern; N = 3, n= 118-146 cells, ***p < 0.001 in student t-test; error bars indicate s.e.m. (**D**) Quantification of the CEP164 localization patterns in 2:2 and 1:1 metaphase hTertRPE1 eGFP-centrin1 cells. N = 3, n = 78 – 107 cells. (**E**) Immunofluorescence images of metaphase hTertRPE1 eGFP-centrin1 stained with DAPI and CEP164 antibodies and treated either with DMSO or centrinone for 18h. (**F**) Immunofluorescence image of an hTert-RPE1-eGFP-centrin1 G2 cell stained with antibodies against CEP164 and centrobin. All scale bars = 5 μm.

### PLK1 controls centrobin localization at centrosomes

In *Drosophila melanogaster* neuroblasts, the PLK1 kinase ortholog Polo controls the specific enrichment of centrobin on the younger centrosome (Gallaud et al., 2020; Januschke et al., 2013). We therefore tested whether mitotic phosphorylation also controls the removal of human centrobin in late prometaphase. Specifically, we tested the contribution of PLK1 and Aurora-A, another key mitotic kinase controlling the centrosome cycle (Barr and Gergely, 2007). RPE eGFP-centrin1 cells were treated for 2 hours with a either mock control (DMSO), 25 or 50 nM of the PLK1 inhibitor BI2536 (Lénárt et al., 2007), or 100 nM of the Aurora-A inhibitor Alisertib (Görgün et al., 2010). Most control-treated cells (67±2%) displayed centrobin on the two daughter centrioles, while 32±2% of the cells displayed three centrobin-positive centrioles (Fig. 4A and B). PLK1 inhibition increased the proportion of cells displaying a third centrobin-positive centriole to respectively 52±3% and 55±1% and resulted in 10% of the cells with four centrobin-positive centrioles (Fig. 4A and B). After Aurora-A inhibition, only 29±5% of the cells harboured centrobin at the two daughter centrioles, 36±5% on three centrioles, and 35±3% on all four centrioles (Fig 4C and D). This indicated that centrobin removal from older centrioles depends both on PLK1 and Aurora-A.

**Figure 4:**
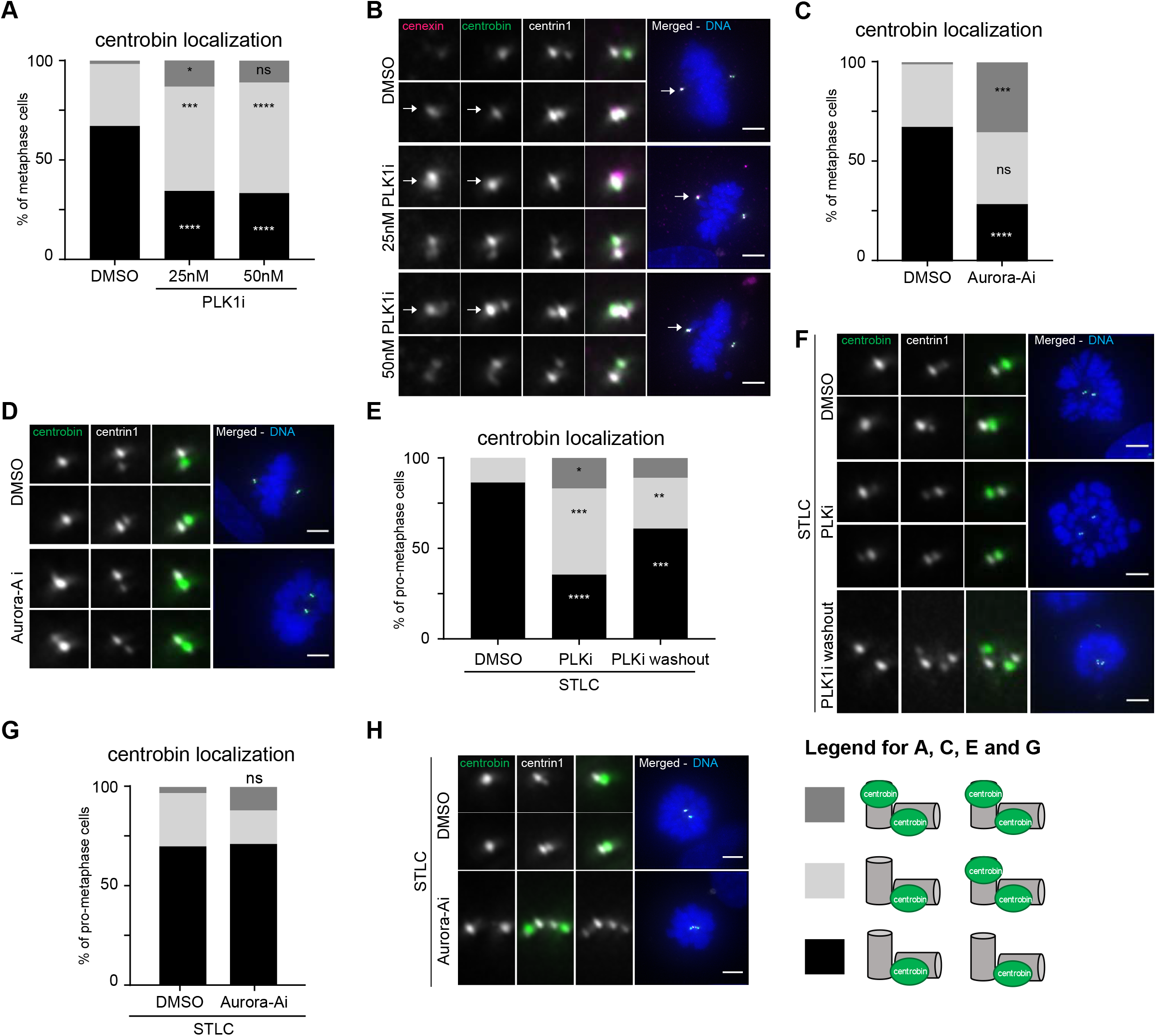
Centrobin removal from mother centrioles depends on Plk1 activity. (**A**) Quantification of centrobin localization patterns in hTertRPE1 eGFP-centrin1 cells treated with DMSO, 25nM or 50 nM of the PLK1 inhibitor BI 2536 for 2h. N = 3; n = 110 – 125 cells; *p<0.05, ***p<0.001, ****p<0.0001 in two-way Anova. (**B**) Immunofluorescence images of metaphase hTertRPE1 eGFP-centrin1 cells treated with DMSO, 25nM or 50 nM of the PLK1 inhibitor BI 2536 for 2h and stained with DAPI and antibodies against cenexin and centrobin. White arrows label the old centrosome. (**C**) Quantification of centrobin localization patterns in hTertRPE1 eGFP-centrin1 cells treated with DMSO, or the Aurora -A inhibitor Alisertib for 2h. N = 3; n = 87 – 110 cells; ***p<0.001, ****p<0.0001 in two-way Anova. (**D**) Immunofluorescent images of metaphase hTertRPE1 eGFP-centrin1 cells treated with DMSO or 100nM Alisertib for 2h and stained with DAPI and an antibody against centrobin. (**E**) Quantification of the centrobin localization patterns in hTertRPE1 eGFP-centrin1 cells arrested in prometaphase with STLC, treated either with DMSO for 4h, with DMSO for 2h and followed by 50nM BI 2536 for 2h (PLK1i), or with BI2536 for 2h followed by DMSO for 2h (PLKi Washout). N = 3, n = 127 – 135 cells. *p<0.05, p<0.01, ***p<0.001, ****p<0.0001 in two-way Anova. (**F**) Immunofluorescence images of prometaphase hTertRPE1 eGFP-centrin1 cells treated as described in (E) and stained with DAPI and an antibody against centrobin. (**G**) Quantification of the centrobin localization patterns in hTertRPE1 eGFP-centrin1 cells arrested in prometaphase with STLC, treated either with DMSO for 4h, or with Alisertib for 4h. N = 3, n = 87 – 140 cells. (**H**) Immunofluorescence images of prometaphase hTertRPE1 eGFP-centrin1 cells treated as described in (G) and stained with DAPI and an antibody against centrobin. All scale bars = 5 μm.

Since PLK1 is activated in G2 by Aurora-A to promote mitotic entry and centrosome separation (Seki et al., 2008), we next aimed to inhibit these two kinases only once cells had entered mitosis. We first arrested the cells in mitosis with STLC for 4 hours, before inhibiting either PLK1 or Aurora-A for 2 hours. STLC treatment and the resulting monopolar spindles did not affect centrobin localization, since we found the same pattern as in untreated cells: most mother centrioles had lost centrobin (Fig 4E and F). PLK1 inhibition led to the rebinding of centrobin on older centrioles in STLC-treated cells, while Aurora-A inhibition had no effect (Fig. 4E-H). This suggested: 1) that Aurora-A does not affect centrobin localization once PLK1 is fully active; 2) that PLK1 is the primary kinase controlling centrobin; 3) that Plk1 inhibition can bring centrobin back to mother centrioles that had already lost it. To confirm the dynamicity of this regulation, we washed out BI2536 in STLC-treated cells after 2 hours and replaced it either with DMSO or again BI2536. Washing out PLK1 inhibition restored normal centrobin localization on two centrioles, confirming the dynamic nature of this regulation in mitosis (Fig. 4E and F).

### PLK1 regulates distal appendage formation independently of centrobin

Since PLK1 also promotes distal appendage formation (Kong et al., 2014), we next investigated the epistasis of this centrosome cycle step in relationship to centrobin removal. Normally, CEP164 is present on both the grandmother and mother centriole in >90% of mitotic cells (Fig. 5A and B). Inhibition of PLK1 for 2 hours in STLC treated cells did not disrupt CEP164 localization, if at all it led to higher CEP164 levels at both centrioles (Fig. 5A and B). In contrast, PLK1 inhibition for 24 hours resulted in 69±2% of cells having CEP164 only at one centriole (Fig. 5C and D), consistent with previous studies (Kong et al., 2014). In vast majority of cells, the CEP164-positive centriole was the grandmother centriole, identified by the higher intensity of the centrin1 signal (Fig. 5E; Tan et al., 2015). This suggested that PLK1 activity is required for the recruitment of CEP164 on the mother centriole prior to mitosis, but not for its maintenance during mitosis. Immunofluorescence analysis of cells treated with BI2536 for 24 hours also indicated that centrobin was more frequently retained on the mother centriole (Fig. 5D and F). Nevertheless, OFD1 was present in 95±1% of cells on both spindle poles (vs. 91±1% in control-treated cells; Fig. 5G and H). This confirmed that centrobin removal is not required for the loading of OFD1 on mother centriole and suggested that PLK1 controls CEP164 recruitment further downstream in the formation of distal appendages. To directly test whether centrobin affects CEP164 recruitment in PLK1-inhibited cells, we combined PLK1 inhibition with the siRNA-mediated depletion of centrobin. Although centrobin was efficiently depleted (Fig. 5I and J), its absence did not rescue the recruitment of CEP164 at the mother centriole: whether centrobin was present or not, PLK1 inhibition led to cells with only one CEP164-positive centriole in 64% of the cases (Fig. 5J and K). We conclude the PLK1 regulates the recruitment of CEP164 and the removal of centrobin via separate pathways.

**Figure 5:**
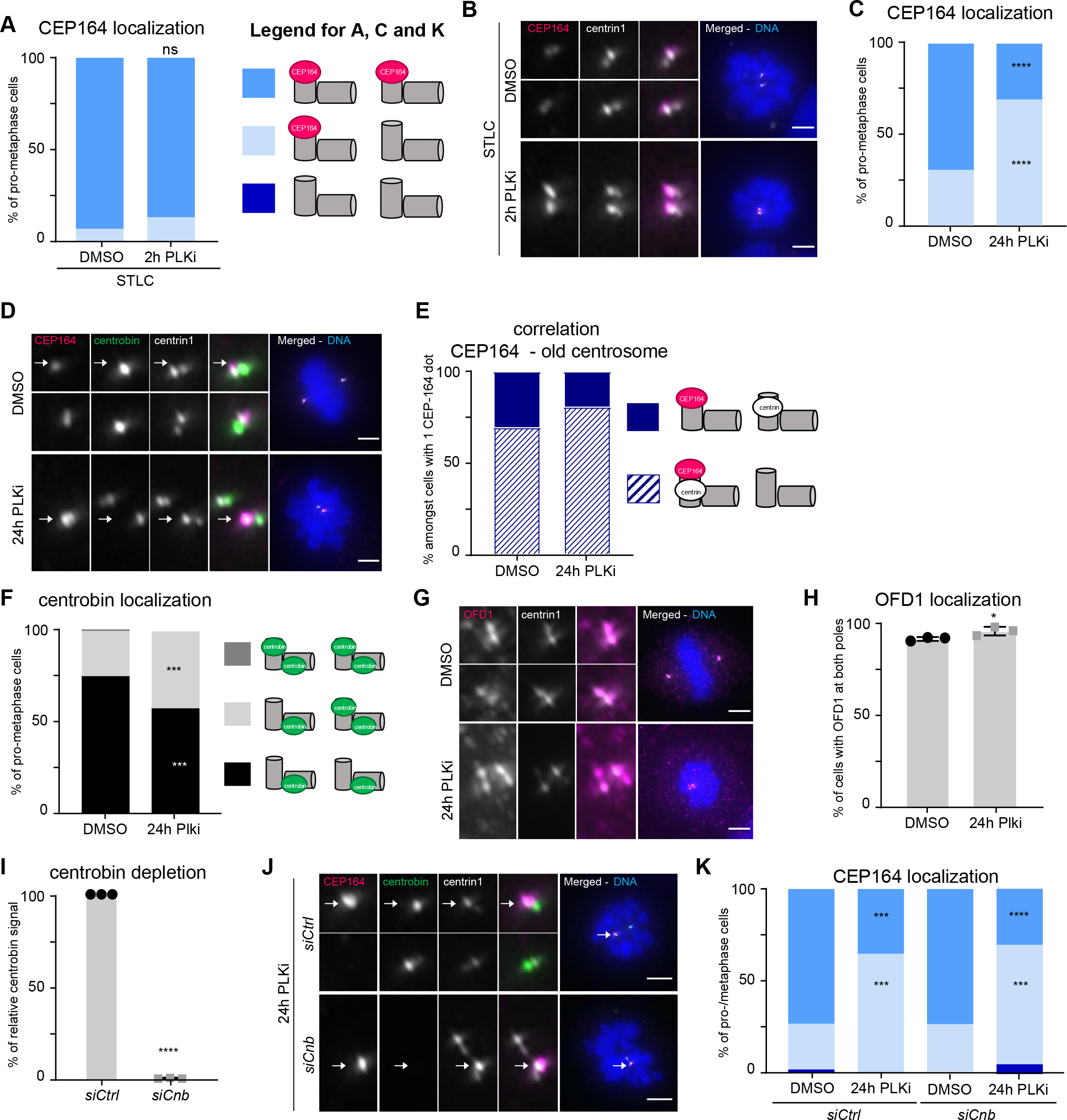
Plk1 controls distal appendage formation and Centrobin removal via independent pathways. (**A**) Quantification of CEP164 localization patterns in hTertRPE1 eGFP-centrin1 cells treated with STLC in combination with 2hrs DMSO or PLK1 inhibition with 50nM BI 2356; N = 3, n = 108 – 127 cells. (**B**). Immunofluorescence images of hTertRPE1 eGFP-centrin1 cells treated with STLC in combination with 2 hrs DMSO or PLK1 inhibition stained with DAPI and CEP164 antibodies. (**C**) Quantification of CEP164 localization patterns in hTertRPE1 eGFP-centrin1 cells treated with DMSO or 24hrs PLK1 inhibition. ****p<0.0001 in student t-test; N = 3, n = 113 – 143 cells. (**D**) Immunofluorescence images of hTertRPE1 eGFP-centrin1 cells treated with DMSO or 24 hrs PLK1 inhibition stained with DAPI and CEP164 antibodies. White arrows identify the old centrosome. (**E**) Correlation between single CEP164 presence and centrosome age based on eGFP-Centrin1 signal. N = 3, n = 29 – 50 cells. (**F**) Quantification of centrobin localization patterns in hTertRPE1 eGFP-centrin1 cells treated with DMSO or 24hrs PLK1 inhibition. ***p<0.001 in student t-test N = 3, n = 113 – 143 cells. (G) Immunofluorescence images of hTertRPE1 eGFP-centrin1 cells treated with DMSO or 24 hrs PLK1 inhibition stained with DAPI and OFD1 antibodies (**H**) Quantification of OFD1 localization patterns in hTertRPE1 eGFP-centrin1 cells treated with DMSO or 24hrs PLK1 inhibition N = 3, n= 108-123 cells; error bars indicate s.e.m. (**I**) Quantification of centrobin depletion efficiency. ****p<0.0001 in student t-test; N = 3, n=124-140 cells; error bars indicate s.e.m. (**J**) Immunofluorescence images of PLK1-inhibited (24hrs) hTertRPE1 eGFP-centrin1 cells treated with control- or centrobin siRNA, stained with DAPI and antibodies against CEP164 and centrobin. White arrows identify the old centrosome. (**K**) Quantification of CEP164 localization patterns in hTertRPE1 eGFP-centrin1 cells treated with indicated treatments. N = 3, n = 108-126 cells. ***p<0.001, ****p<0.0001 in two-way Anova comparing DMSO and PLK1-inhibition. All scale bars = 5 μm

### Cenexin regulates recruitment of CEP164 and centrobin removal through separate pathways

Finally, we investigated the role of the core sub-distal appendage protein cenexin in the regulation of the centrobin and the CEP164 pathways. Cenexin has been reported to modulate PLK1 activity at centrosomes (Colicino et al., 2019); moreover, in mouse embryonic carcinoma cells F9, cenexin is required for the formation of both distal and sub-distal appendages (Ishikawa et al., 2005; Tateishi et al., 2013). In RPE cells, however, it was reported to be only required for the formation of sub-distal appendages (Kuhns et al., 2013; Tanos et al., 2013). While using two different siRNAs against cenexin to rule out any off-target effect, we found that efficient cenexin depletion (>80%, see Supplementary Fig. 2A and B) led in most cells to the same configuration as the one seen after PLK1 inhibition: 1) only the grandmother centriole retained CEP164 (Fig. 6A and B, Supplementary Fig. 2C; note that grandmother centrioles were identified based on the highest intensity of the eGFP-centrin1 signal); 2) centrobin was present on the mother centriole (Fig. 6C and D, Supplementary Fig. 2D); and 3) OFD1 recruitment was not affected (Fig. 6E-F). This indicated that, as PLK1, cenexin is required for the removal of centrobin and the recruitment of CEP164 at the mother centriole, but that it does not affect OFD1 localization. To test for the epistasis of CEP164 recruitment and centrobin removal, we co-depleted both cenexin and centrobin and found that centrobin depletion did not rescue CEP164 binding to the mother centriole in cenexin-depleted cells (Fig. 6G and H; note that the co-depletions were as efficient as the single depletions; Sup Fig 2E-H). We conclude that cenexin regulates centrobin and CEP164 separately, and in an OFD1-independent manner.

**Figure 6:**
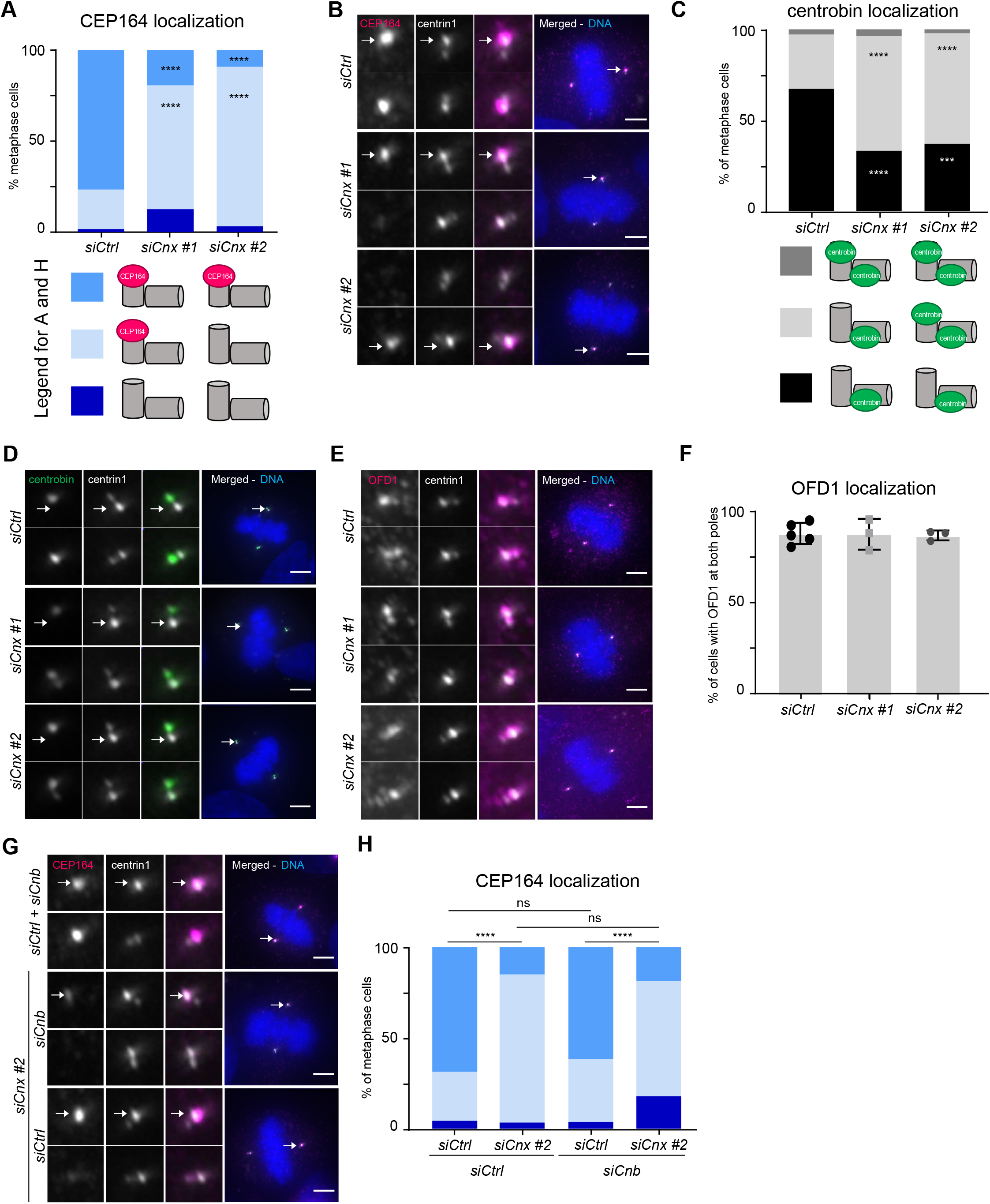
Cenexin controls centrobin removal and subdistal appendage formation independently. (**A**) Quantification of CEP164 localization patterns in hTertRPE1 eGFP-centrin1 cells treated with either control- or two different cenexin siRNAs, ****p<0.0001 in two-way Anova, N = 3, n = 123-160 cells. (**B**) Immunofluorescence images of hTertRPE1 eGFP-centrin1 cells treated with indicated siRNAs stained with DAPI and CEP164 antibodies. (**C**) Quantification of centrobin localization patterns in hTertRPE1 eGFP-centrin1 cells treated with indicated siRNAs; ****p<0.0001 in two-way Anova; N ≥ 3, n = 124 – 160 cells. (**D** and **E**) Immunofluorescence images of hTertRPE1 eGFP-centrin1 cells treated with indicated siRNAs stained with DAPI and (D) centrobin or (E) OFD1 antibodies. (**F**) Quantification of OFD1 localization patterns in hTertRPE1 eGFP-centrin1 cells treated with indicated siRNAs; N = 3, n = 119 – 194 cells; error bars indicate s.e.m. (**G**) Immunofluorescence images of hTertRPE1 eGFP-centrin1 cells treated with indicated siRNAs, stained with DAPI and CEP164 antibodies. White arrows identify the old centrosome. (**H**) Quantification of CEP164 localization patterns in hTertRPE1 eGFP-centrin1 cells treated with indicated siRNAs; ****p<0.0001 in two-way Anova; N = 3, n = 134-137 cells. All scale bar = 5 μm

We next extended our dependency analysis to the centrosomal proteins CEP128 and Centriolin, which act downstream of cenexin in terms of building up sub-distal appendages (Mazo et al., 2016). Using the published hTert-RPE1 cenexin, CEP128 or centriolin CRISPR/Cas9 knock-out cell lines, we first found that only cenexin deletion partially prevented the recruitment of CEP164, and that no defects could be seen in CEP128 or centriolin knock-out cells (Fig. 7A and B; note that the percentage of cells with only one CEP164-positive centriole was lower compared to cenexin-depleted cells, see discussion). Second, our analysis indicated that centrobin was present on all four centrioles in all three knock-out cell lines, but that OFD1 localization was unchanged (Fig. 7C-F). This confirmed that the recruitment of distal appendage proteins does not require the removal of centrobin from older centrioles. Third, we conclude the removal of centrobin from the maturing centrioles requires the presence of sub-distal appendage proteins. In contrast, distal appendages, appear not to be required, since the depletion of the distal appendage protein CEP164 had no effect of centrobin localization (Supplementary Fig. 3).

**Figure 7:**
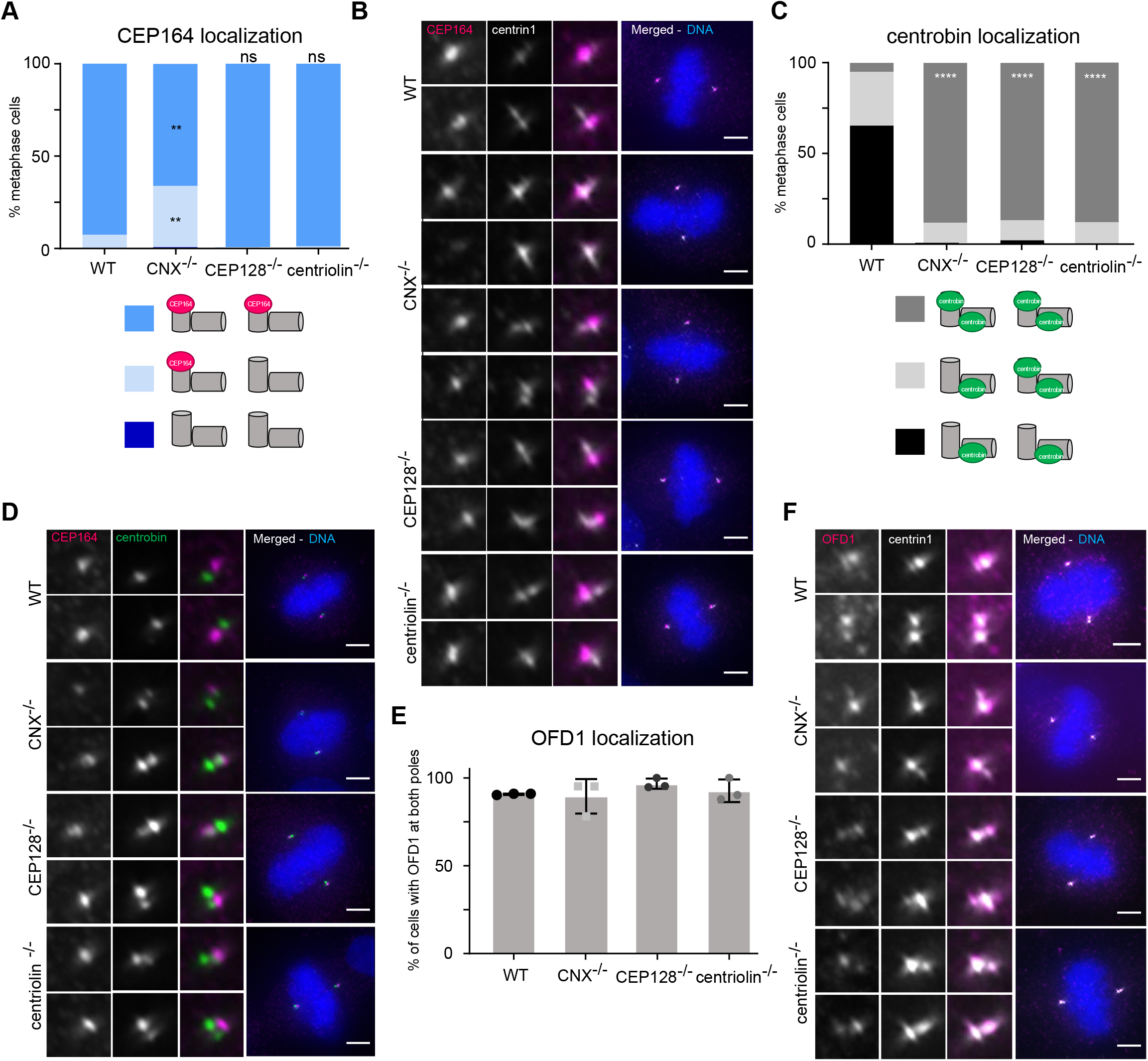
Sub-distal appendages are required for centrobin removal, but not for distal appendage assembly. (**A**) Quantification of CEP164 localization patterns in the WT hTertRPE1 cell line and cenexin, CEP128 or centriolin knockouts; **p<0.01 in two-way Anova N = 3, n = 119-155 cells. (**B**) Immunofluorescence images of WT or respective knock-out metaphase hTert-RPE1 cells stained with DAPI and antibodies against CEP164 and centrin1. (**C**) Quantification of the centrobin localization patterns in the indicated hTertRPE1 cells; ****p<0.0001 in two-way Anova, N = 3, n = 119 – 154 cells. (**D**) Immunofluorescence images of indicated hTertRPE1 cells stained with DAPI and antibodies against CEP164 and centrobin. (**E**) Quantification of OFD1 localization patterns in indicated hTert-RPE1 cell lines; N = 3, n = 115 – 129 cells; error bars indicate s.e.m. (**F**) Immunofluorescence images of indicated hTert-RPE1 cells stained with DAPI and antibodies against OFD1 and centrin1. All scale bars = 5 μm.

## Discussion

In this present study we investigated how the localization of the daughter-centriole specific protein centrobin is controlled over the cell cycle in human cells, and whether centrobin acts as a placeholder protein for distal appendage proteins on the mother centriole. Our results imply that centrobin is removed from mother centriole in a dynamic manner and transferred to higher affinity sites on daughter centrioles once cells reach mitosis. Centrobin removal requires the activity of the mitotic kinase Plk1 and the presence of sub-distal appendage proteins on the mature centrioles. Finally, our results imply that centrobin does not act as a placeholder for the distal appendage proteins, but rather that centrobin removal and distal appendage formation are promoted by Plk1 via separate pathways.

Previous work in *Drosophila melanogaster* had shown that centrobin is transferred in a Plk1-dependent manner within the younger centrosome from the mother centriole to the daughter centriole in metaphase (note that in flies centrobin is present not on daughter centrioles but on the younger centrosome (Januschke et al., 2013)); moreover, a later study also speculated that this transfer might be direct (Gallaud et al., 2020). Here, we show that removal from the mother centriole until metaphase is dynamic, as centrobin-pools can co-exist on mother and daughter-centrioles in G2, prophase and early prometaphase cells; moreover, we demonstrate that the presence of at least one daughter centriole is essential to remove centrobin from the mother centriole. This implies the existence of higher-affinity binding sites on the daughter centrioles that strip the mother-centriole bound pool of centrobin.

While our data indicate that centrobin removal is regulated in a reversible manner via Plk1 activity, it is unclear at which level this kinase acts. In theory it could decrease the affinity of the centrobin-binding site on the mother centriole, increase it on the daughter centrioles, or switch the binding preference by directly phosphorylating centrobin itself, in line with the fact that centrobin is a Plk1 substrate (Lee et al., 2010). The subdistal appendage proteins cenexin, CEP128 and centriolin, which in metaphase are present exclusively at the grandmother centriole, are also required for centrobin removal from the mother centriole. Given the vast distances between the two spindle poles at this stage, this control is most likely indirect, although one cannot completely exclude that subdistal appendages on the grandmother centriole may affect the mother centriole in the preceding interphase. One likely pathway is that the subdistal appendages regulate Plk1 activity, consistent with previous studies showing that the presence of cenexin enhances Plk1 activity (Colicino et al., 2019); whether CEP128 and centriolin play a similar role, remains to be investigated.

Finally, our results indicate that centrobin does not act as a placeholder for distal appendage proteins, in contrast to a previously proposed model (Wang et al., 2018). Our data identify multiple experimental conditions, in which centrobin presence at mother centriole does not prevent the recruitment of OFD1, which is recruited early, and of CEP164, which is recruited last during distal appendage formation. This co-existence can even be seen in wild-type cells, as CEP164 is recruited to the mother centriole in late G2, whilst most centrobin is removed during prometaphase/metaphase. Our investigation of subdistal appendage proteins further indicates that cenexin is partially required for the loading of CEP164, but not of OFD1. This role is specific for cenexin, as knockouts of CEP128 and centriolin did not reveal any changes in CEP164 localization, confirming that centrobin removal and CEP164 recruitment are independent. Finally, our data confirm that distal appendage protein formation requires Plk1 activity, consistent with previous studies (Kong et al., 2014; Tanos et al., 2013). This dependency is, however, only transient, since Plk1 inhibition does not affect the maintenance of distal appendages at later stages. Overall, we postulate that Plk1 controls centriole maturation via separate pathways, one involving centrobin removal, the second controlling distal appendage formation. This raises the question of the reasons for centrobin removal, since its presence does not impact distal appendage formation. In *Drosophila*, centrobin localization has to be controlled since it regulates the fate of the young centrosome, enabling it to organize microtubules and to be retained by the stem cell during asymmetric cell division (Januschke et al., 2013). In mammalian cells, however, the reasons for controlling centrobin still remain to be uncovered. Our characterization of 1:1 cells indicate that the presence of centrobin at the mother centriole on one pole does neither affect mitotic progression, nor change the incidence of chromosome segregation errors, indicating that its removal is not acutely necessary for centrosome/spindle pole function in mitosis. Rather we speculate that timely removal of centrobin in (pro-)metaphase might be linked to the faithfulness of the centrosome duplication cycle and to cilia formation, since it has been shown that overexpressing centrobin leads to overelongated centrioles and may affect the axoneme structure during ciliogenesis (Gudi et al., 2015; Ogungbenro et al., 2018).

## Supplementary figures legends

**Supplementary Figure 1:**
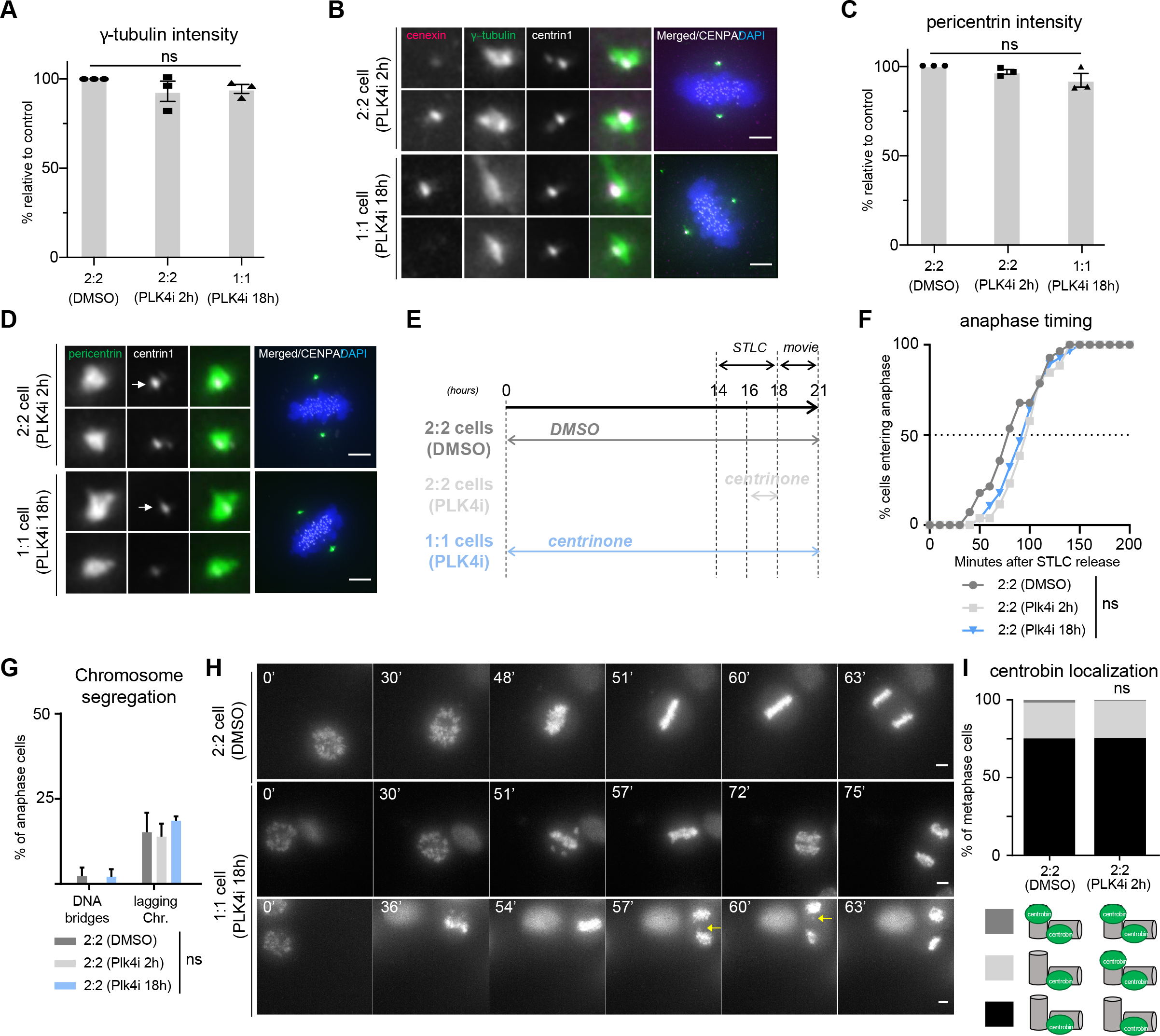
hTert-RPE1 1:1 cells display normal centrosome function and chromosome segregation. (**A**) Quantification of relative γ-tubulin levels at centrosomes in cells treated with DMSO or 250nM centrinone for 2h or 18h. N = 3, n = 114-139 cells; error bars indicate s.e.m. (**B**) Immunofluorescence of metaphase hTert-RPE1 cells treated with centrinone for 2h or 18h stained with DAPI and antibodies against cenexin and γ-tubulin. (**C**) Quantification of relative pericentrin levels at centrosomes in cells treated with DMSO, 2h centrinone, or 18h centrinone. N = 3, n = 93-194 cells; error bars indicate s.e.m. (**D**) Immunofluorescence of metaphase hTert-RPE1 cells treated with centrinone for 2h or 18h stained with DAPI and antibodies against cenexin and γ-tubulin. (**E**) Experimental design for the STLC release experiment: cells were treated with DMSO or 250 nM centrinone for 2h or 18h, then blocked in a monopolar conformation with 5μM STLC for 4h, before washing out STLC and observing mitotic progression with SiR-DNA in live cell movies. (**F**) Cumulative frequency of anaphase entry after STLC release (t = 0) in indicated cell lines; N = 4, n = 34 - 38 cells. (**G**) Anaphase outcomes of indicated cells after STLC release; N = 4, n = 16-38 cells. (**H**) Time-lapse image sequences of hTert-RPE1 cells treated with DMSO or 18h centrinone after STLC release. Yellow arrows indicate lagging chromosomes. Timing is in mins using NEBD as T = 0. (**I**) Quantification of the centrobin localization patterns in hTertRPE1 eGFP-Centrin1/CENPA-eGFP cells treated with DMSO or 250nM centrinone for 2h. N = 3, n = 143 - 151 cells. All scale bars = 5 μm.

**Supplementary Figure 2:**
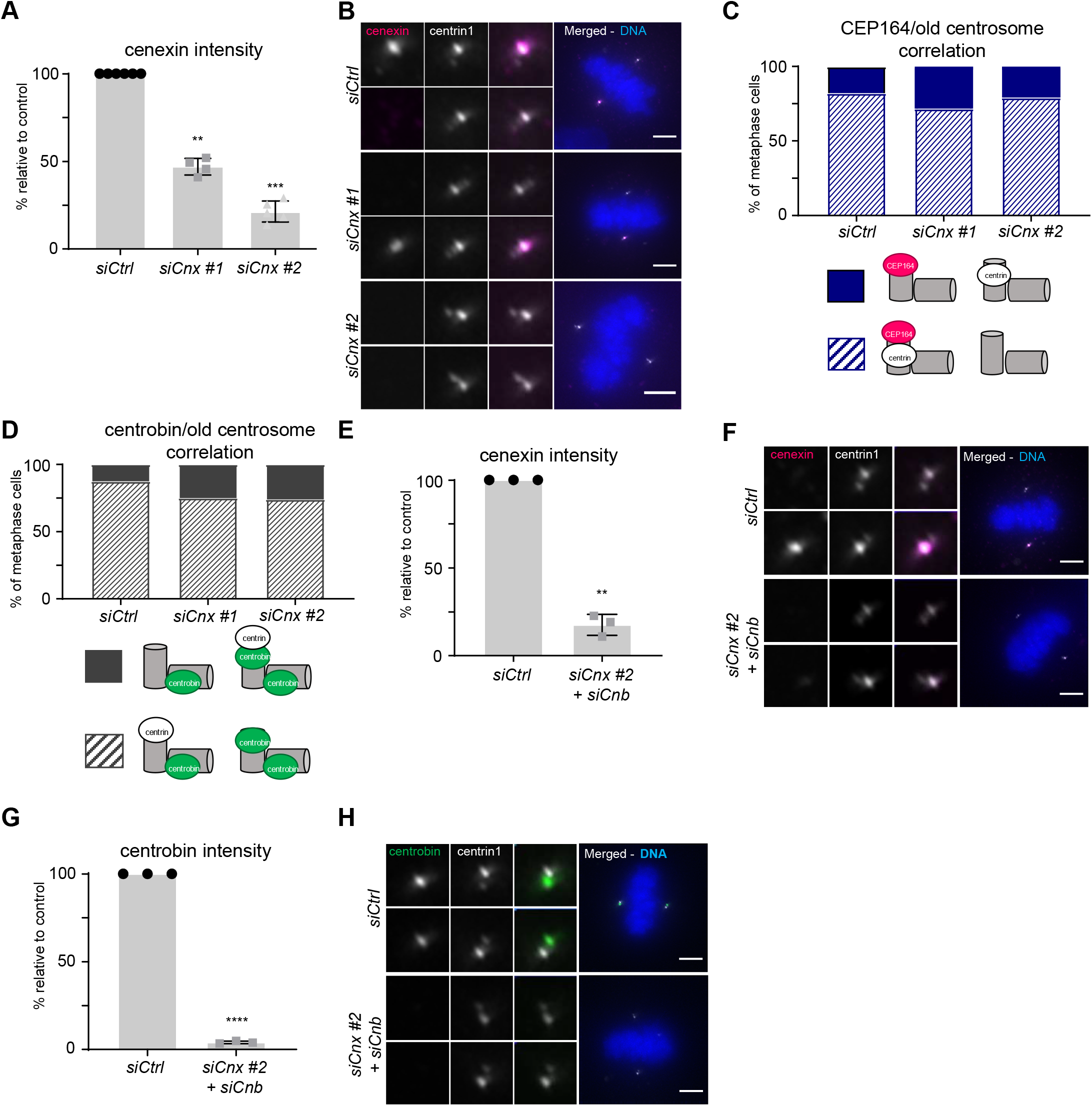
Cenexin and centrobin depletion efficiencies. (**A**) Quantification of cenexin levels in hTertRPE1 eGFPcentrin1 cells treated with *siCtrl*, *siCnx #1* or *siCnx #2* for 48h; **p<0.01 in Anova test; N ≥ 4, n = 153-231 cells. (**B**) Immunofluorescence images of metaphase hTertRPE1 eGFPcentrin1 cells treated with *siCtrl*, *siCnx #1* or *siCnx #2* and stained with DAPI and cenexin antibodies. (**C** and **D**) Correlation between the presence of CEP164 (C) or centrobin (D) at the centriole with the highest eGFP-centrin1 signal; N ≥ 3, n = 31-95 cells. (**E**) Quantification of cenexin levels in double depleted cenexin/centrobin hTertRPE1 eGFP-centrin1 cells; **p<0.01 in t-test; N = 3, n = 116 – 127 cells. (**F**) Representative images of cells quantified in (E). (**G**) Quantification of centrobin levels in double depleted cenexin/centrobin hTertRPE1 eGFP-centrin1 cells; ****p<0.0011 in t-test; N = 3, n = 136 – 144 cells. (**H**) Representative images of cells quantified in (G). All scale bars = 5 μm.

**Supplementary Figure 3:**
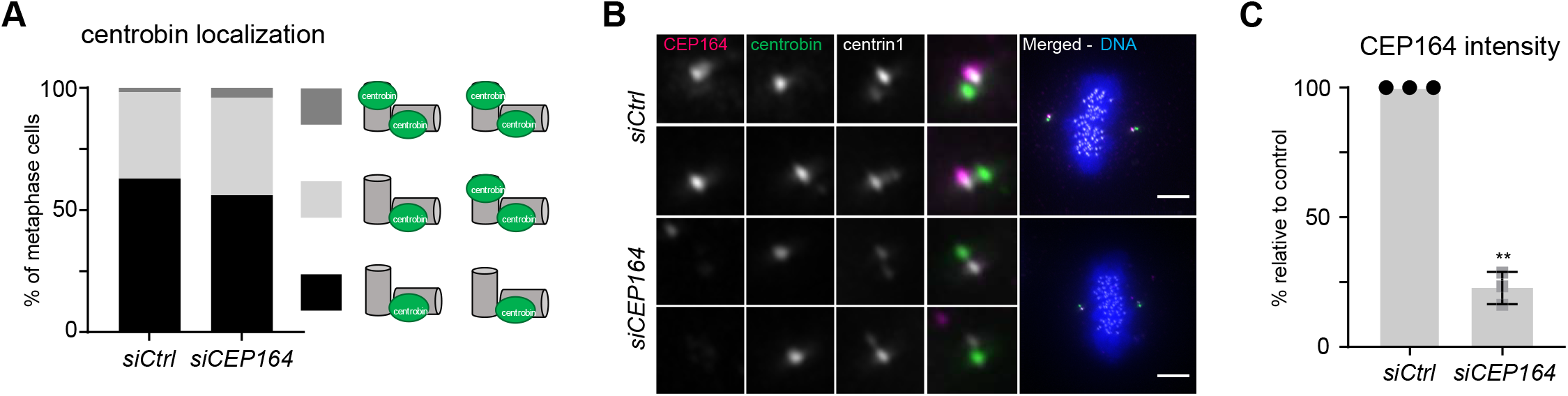
The distal appendage protein CEP164 does not affect centrobin localization. (**A**) Quantification of the centrobin localization patterns in hTertRPE1 eGFP-centrin1/CENPA-GFP cells treated with Ctrl or CEP164 siRNAs; N = 3, n = 91 – 158 cells. (**B**) Immunofluorescence images of metaphase hTertRPE1 eGFP-centrin1/CENPA-GFP cells stained with DAPI and antibodies against CEP164 and centrobin. Scale bar = 5 μm. (**C**) Quantification of CEP164 levels in hTertRPE1 eGFP-centrin1/CENPA-GFP cells; N = 3, n = 91-158 cells, error bars indicate s.e.m.

## Methods

### Cell culture and drug treatments

hTERT-RPE1 cells, hTERT-RPE1 eGFP-centrin1, hTERT-RPE1 eGFP-Centrin1/eGFP-CENP-A (both gifts of A. Khodjakov (Magidson et al., 2011)), hTERT-RPE1 Cenexin KO, Cep128 KO, Centriolin KO and their parental cell line (gift of B. Tsou; Mazo et al., 2016), were all grown in DMEM (Thermofisher, Switzerland), supplemented with 10% FCS, 100 U/ml penicillin, and 100 mg/ml streptomycin (all Life Technologies, Switzerland) at 37°C and 5% CO_2_. For live-cell imaging, cells were kept in their normal medium if a CO_2_ chamber was used or otherwise cultured in Leibovitz’s L-15 medium, no phenol red (Thermofisher), supplemented with 5% FCS. To inhibit PLK1, 25 nM or 50 nM BI 2536 (Axon Lab AG, Switzerland) was used, for either 2h or 24h. To inhibit Aurora-A, 100 nM Alisertib (Selleckchem, USA) was used for 2h. For the PLK1 inhibition washout experiments, cells were treated with 5 μM STLC for 4h to maintain them in prometaphase, 50 nM BI 2536 was added for 2h before adding back medium containing only STLC for 2h. To obtain cells with only one centriole at each pole, cells were treated with 250 nM centrinone (PLK4 inhibitor; Tocris, UK) for 18h. Alternatively, a short 2h treatment was added as a negative control.

### SiRNA-mediated protein depletions

In general siRNA treatments were performed for 48 hours applying 40 nmol siRNAs and 1% Lipofectamine RNAimax (Invitrogen) using manufacturers protocol. To deplete cenexin, we applied 180 nmol siRNA and 3% Lipofectamine RNAimax (Invitrogen) instead; to prevent centriole duplication and obtain 1:1 cells, Sas-6 was depleted for only 24 hours. The target sequences of the siRNAs used are: *siCEP164* (mix of 5’ – CAGGTGACATTTACTATTTCA 3’, and 5’ – ACCACTGGGAATAGAAGACAA – 3’), *siCnb* (mix of 5’ – TGGAAATGGCAGAACGAGA – 3’, 5’- GCATGAGGCTGAGCGGACA – 3’, 5’ – GCCCAAGAATTGAGTCGAA – 3’, and 5’ – CTCCAAACCTCACGTGATA – 3’), *siCnx #1* (5’- TGGCTGAGACTGAGCACGA – 3’), *siCnx #2* (5’ – GGCACAACATCGAGCGCAT 3’), siCtrl (5’-AAGGACCTGGAGGTCTGCTGT – 3’) and *siSas-6* (5’ – GCACGTTAATCAGCTACAA – 3’)

### Immunofluorescence

Cells were grown on acid-etched glass coverslips and either fixed for 6 min at −20°C with ice-old methanol, or for 15min with a formaldehyde fixative (0.05M PIPES, 0.01M EGTA, 1mM MgCl_2_, 0,2% Triton X-100, 4% Formaldehyde) at RT. Coverslips were washed with Phosphate-buffered saline solution (PBS) three times before adding a blocking buffer for 1h at RT (7.5% Bovine Serum Albumin, 0.25% sodium azide in PBS). Primary antibodies were diluted in blocking buffer and added for 1h at RT followed by 3 PBS washes. The following primary antibodies were used: mouse anti-cenexin (1:1000, Abcam, UK, ab43840), mouse anti-centrin1 (1:2000, clone 20H5 Merck Millipore, Switzerland), mouse anti-centrobin (1:1000, Abcam ab70448), rabbit anti-CEP164 (1:1000, gift of E. Nigg; Graser et al., 2007), rabbit anti-OFD1 (1:1000, Abcam ab222837), mouse anti-pericentrin (1:1000, Abcam ab28144), rabbit anti-γ-tubulin (1:2000; Wilhelm et al., 2019). Cross-absorbed secondary antibodies tagged with Alexa-fluorophores were diluted in blocking buffer (1:400, Invitrogen, Switzerland) and applied for 30 min at RT. VECTASHIELD with DAPI (Vector Laboratories, Switzerland) was used to mount the coverslips, and images were taken with a wide-field Olympus Deltavision (GE Healthcare, USA), with a 60x 1.4NA oil objective and a DAPI/FITC/TRITC/Cy5 (Chroma, USA) filter set. The z-stack images were recorded in 0.2 μm steps with a Coolsnap HQ2 CCD camera (Roper Scientific) and the Softworx software (GE Healthcare). Image stacks were deconvolved and protein levels at centrioles were quantified based on maximal intensity projections using Softworx Explorer (GE Healthcare). To quantify pericentriolar protein we applied with Imaris (Bitplane, Switzerland) a 0.5 μm diameter sphere centred on the eGFP-centrin1 signal and quantified the summed intensity within this sphere.

### Readout for centrosome age

In most of the experiments, cenexin signal was used to identify the grandmother centriole, and therefore which centrosome is the old one. For experiments where cenexin was depleted and for experiments done with RPE cells knocked-out for cenexin, CEP128 or centriolin, centrin1 signal was used as a readout (Gasic et al., 2015), and only cells with a difference higher than 5% in centrin1 signal between the two centrosomes were analysed. This difference was calculated with the following formula: 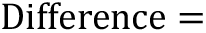 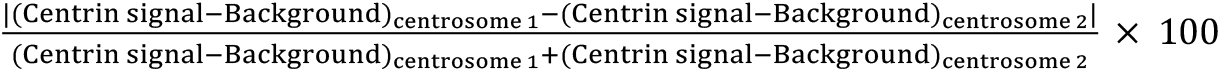. Centrin was also used to distinguish between old and young centrosome in experiments where CEP164 localization was studied.

### Statistical analysis

If not indicated differently, the data is presented as the mean of multiple independent experiments and error bars represent the standard error of the mean, unless otherwise specified. Significance was determined using Graphpad Prism®, and were represented presented in this work according to the following codification: *p<0.05, **p<0.01, ***p<0.001 and ****p<0.0001. An unpaired two-tailed t test assessed the depletion efficiency for cenexin, Centrobin, or CEP164. For all the data describing the changes in centrobin and CEP164 localization, a two-way ANOVA with Tukey’s multiple comparison test was chosen. All illustrations and graphs were created using ImageJ Fiji, Graphpad Prism and Adobe Illustrator.

## Acknowledgments

Authors are grateful to A. McAinsh (University of Warwick, UK), A. Khodjakov (NY State University, USA), B. Tsou (Cornell University, NY, USA) for cell lines, E. Nigg (University of Basel, Switzerland) for CEP164 antibodies, Monica Gotta (University of Geneva, Switzerland) and members of the Meraldi laboratory for helpful discussions and support, especially E. Galster for her help in cell lines’ maintenance and A.-M. Olziersky, L. Romanens, D. Dudka, T. Whilhelm and D. Dwivedi for critical comments on the manuscript. This work was supported by the Swiss National Science Foundation (SNF) project grant (No. 31003A_179413) and the University of Geneva.

## Author contributions

The project was initiated by M.L.R.B. and P.M. and directed by P.M. M.L.R.B. and D.H. performed all the experiments. M.L.R.B. analysed the results and M.L.R.B. and P.M. interpreted the data and wrote the manuscript.

## References

Barr, A. R. and Gergely, F. (2007). Aurora-A: the maker and breaker of spindle poles. J Cell Sci 120, 2987–2996.

Bornens, M. (2002). Centrosome composition and microtubule anchoring mechanisms. Curr Opin Cell Biol 14, 25–34.

Bowler, M., Kong, D., Sun, S., Nanjundappa, R., Evans, L., Farmer, V., Holland, A., Mahjoub, M. R., Sui, H. and Loncarek, J. (2019). High-resolution characterization of centriole distal appendage morphology and dynamics by correlative STORM and electron microscopy. Nat Commun 10, 993–15.

Chong, W. M., Wang, W.-J., Lo, C.-H., Chiu, T.-Y., Chang, T.-J., Liu, Y.-P., Tanos, B., Mazo, G., Tsou, M.-F. B., Jane, W.-N., et al. (2020). Super-resolution microscopy reveals coupling between mammalian centriole subdistal appendages and distal appendages. Elife 9.

Colicino, E. G., Stevens, K., Curtis, E., Rathbun, L., Bates, M., Manikas, J., Amack, J., Freshour, J. and Hehnly, H. (2019). Chromosome misalignment is associated with PLK1 activity at cenexin-positive mitotic centrosomes. Mol Biol Cell 30, 1598–1609.

DeBonis, S., Skoufias, D. A., Lebeau, L., Lopez, R., Robin, G., Margolis, R. L., Wade, R. H. and Kozielski, F. (2004). In vitro screening for inhibitors of the human mitotic kinesin Eg5 with antimitotic and antitumor activities. Mol. Cancer Ther. 3, 1079–1090.

Delgehyr, N., Sillibourne, J. and Bornens, M. (2005). Microtubule nucleation and anchoring at the centrosome are independent processes linked by ninein function. J Cell Sci 118, 1565–1575.

Dudka, D., Castrogiovanni, C., Liaudet, N., Vassal, H. and Meraldi, P. (2019). Spindle-Length-Dependent HURP Localization Allows Centrosomes to Control Kinetochore-Fiber Plus-End Dynamics. Curr Biol 29, 3563–3578.e6.

Gallaud, E., Ramdas Nair, A., Horsley, N., Monnard, A., Singh, P., Pham, T. T., Salvador Garcia, D., Ferrand, A. and Cabernard, C. (2020). Dynamic centriolar localization of Polo and Centrobin in early mitosis primes centrosome asymmetry. PLoS Biol 18, e3000762.

Gasic, I., Nerurkar, P. and Meraldi, P. (2015). Centrosome age regulates kinetochore microtubule stability and biases chromosome mis-segregation. Elife 4, e07909.

Gönczy, P. and Hatzopoulos, G. N. (2019). Centriole assembly at a glance. J Cell Sci 132.

Görgün, G., Calabrese, E., Hideshima, T., Ecsedy, J., Perrone, G., Mani, M., Ikeda, H., Bianchi, G., Hu, Y., Cirstea, D., et al. (2010). A novel Aurora-A kinase inhibitor MLN8237 induces cytotoxicity and cell-cycle arrest in multiple myeloma. Blood 115, 5202–5213.

Graser, S., Stierhof, Y.-D., Lavoie, S. B., Gassner, O. S., Lamla, S., Le Clech, M. and Nigg, E. A. (2007). Cep164, a novel centriole appendage protein required for primary cilium formation. 179, 321–330.

Gudi, R., Haycraft, C. J., Bell, P. D., Li, Z. and Vasu, C. (2015). Centrobin-mediated regulation of the centrosomal protein 4.1-associated protein (CPAP) level limits centriole length during elongation stage. J Biol Chem 290, 6890–6902.

Gudi, R., Zou, C., Dhar, J., Gao, Q. and Vasu, C. (2014). Centrobin-centrosomal protein 4.1-associated protein (CPAP) interaction promotes CPAP localization to the centrioles during centriole duplication. J Biol Chem 289, 15166–15178.

Ishikawa, H., Kubo, A., Tsukita, S. and Tsukita, S. (2005). Odf2-deficient mother centrioles lack distal/subdistal appendages and the ability to generate primary cilia. Nat Cell Biol 7, 517–524.

Januschke, J., Reina, J., Llamazares, S., Bertran, T., Rossi, F., Roig, J. and Gonzalez, C. (2013). Centrobin controls mother-daughter centriole asymmetry in Drosophila neuroblasts. Nat Cell Biol 15, 241–248.

Kong, D., Farmer, V., Shukla, A., James, J., Gruskin, R., Kiriyama, S. and Loncarek, J. (2014). Centriole maturation requires regulated Plk1 activity during two consecutive cell cycles. J. Cell Biol. 206, 855–865.

Kuhns, S., Schmidt, K. N., Reymann, J., Gilbert, D. F., Neuner, A., Hub, B., Carvalho, R., Wiedemann, P., Zentgraf, H., Erfle, H., et al. (2013). The microtubule affinity regulating kinase MARK4 promotes axoneme extension during early ciliogenesis. J. Cell Biol. 200, 505–522.

Lange, B. M. and Gull, K. (1995). A molecular marker for centriole maturation in the mammalian cell cycle. 130, 919–927.

Lee, J., Jeong, Y., Jeong, S. and Rhee, K. (2010). Centrobin/NIP2 is a microtubule stabilizer whose activity is enhanced by PLK1 phosphorylation during mitosis. J Biol Chem 285, 25476–25484.

Leidel, S., Delattre, M., Cerutti, L., Baumer, K. and Gönczy, P. (2005). SAS-6 defines a protein family required for centrosome duplication in C. elegans and in human cells. Nat Cell Biol 7, 115–125.

Lénárt, P., Petronczki, M., Steegmaier, M., Di Fiore, B., Lipp, J. J., Hoffmann, M., Rettig, W. J., Kraut, N. and Peters, J.-M. (2007). The small-molecule inhibitor BI 2536 reveals novel insights into mitotic roles of polo-like kinase 1. Curr Biol 17, 304–315.

Loncarek, J. and Bettencourt-Dias, M. (2018). Building the right centriole for each cell type. J. Cell Biol. 217, 823–835.

Magidson, V., O’Connell, C. B., Loncarek, J., Paul, R., Mogilner, A. and Khodjakov, A. L. (2011). The Spatial Arrangement of Chromosomes during Prometaphase Facilitates Spindle Assembly. Cell 146, 555–567.

Mazo, G., Soplop, N., Wang, W.-J., Uryu, K. and Tsou, M.-F. B. (2016). Spatial Control of Primary Ciliogenesis by Subdistal Appendages Alters Sensation-Associated Properties of Cilia. Dev Cell 39, 424–437.

Ogungbenro, Y. A., Tena, T. C., Gaboriau, D., Lalor, P., Dockery, P., Philipp, M. and Morrison, C. G. (2018). Centrobin controls primary ciliogenesis in vertebrates. J. Cell Biol. 217, 1205–1215.

Seki, A., Coppinger, J. A., Jang, C.-Y., Yates, J. R. and Fang, G. (2008). Bora and the kinase Aurora a cooperatively activate the kinase Plk1 and control mitotic entry. Science 320, 1655–1658.

Singla, V., Romaguera-Ros, M., Garcia-Verdugo, J. M. and Reiter, J. F. (2010). Ofd1, a human disease gene, regulates the length and distal structure of centrioles. Dev Cell 18, 410–424.

Tan, C. H., Gasic, I., Huber-Reggi, S. P., Dudka, D., Barisic, M., Maiato, H. and Meraldi, P. (2015). The equatorial position of the metaphase plate ensures symmetric cell divisions. Elife 4, e05124.

Tanos, B. E., Yang, H.-J., Soni, R., Wang, W.-J., Macaluso, F. P., Asara, J. M. and Tsou, M.-F. B. (2013). Centriole distal appendages promote membrane docking, leading to cilia initiation. Genes Dev 27, 163–168.

Tateishi, K., Yamazaki, Y., Nishida, T., Watanabe, S., Kunimoto, K., Ishikawa, H. and Tsukita, S. (2013). Two appendages homologous between basal bodies and centrioles are formed using distinct Odf2 domains. 203, 417–425.

Tischer, J., Carden, S. and Gergely, F. (2021). Accessorizing the centrosome: new insights into centriolar appendages and satellites. Curr. Opin. Struct. Biol. 66, 148–155.

Vásquez-Limeta, A. and Loncarek, J. (2021). Human centrosome organization and function in interphase and mitosis. Semin. Cell Dev. Biol.

Wang, L., Failler, M., Fu, W. and Dynlacht, B. D. (2018). A distal centriolar protein network controls organelle maturation and asymmetry. Nat Commun 9, 3938.

Wilhelm, T., Olziersky, A.-M., Harry, D., De Sousa, F., Vassal, H., Eskat, A. and Meraldi, P. (2019). Mild replication stress causes chromosome mis-segregation via premature centriole disengagement. Nat Commun 10, 3585–14.

Wong, Y. L., Anzola, J. V., Davis, R. L., Yoon, M., Motamedi, A., Kroll, A., Seo, C. P., Hsia, J. E., Kim, S. K., Mitchell, J. W., et al. (2015). Cell biology. Reversible centriole depletion with an inhibitor of Polo-like kinase 4. Science 348, 1155–1160.

Zou, C., Li, J., Bai, Y., Gunning, W. T., Wazer, D. E., Band, V. and Gao, Q. (2005). Centrobin: a novel daughter centriole-associated protein that is required for centriole duplication. 171, 437–445.

